# A Novel Hypothesis for Migraine Disease Mechanism: The Creation of a New Attractor Responsible for Migraine Disease Symptoms

**DOI:** 10.1101/2022.11.13.516319

**Authors:** Farnaz Garehdaghi, Yashar Sarbaz, Elham Baradari

**Affiliations:** Modeling Biological System’s Laboratory, Department of Biomedical Engineering, Faculty of Electrical and Computer Engineering, University of Tabriz, Tabriz, Iran

**Keywords:** Headache, Ictal, Migraineur, Complex Dynamic System, Chaotic Attractor, Chua’s System

## Abstract

Migraine Disease (MD) is one of the primary headaches in which the pathophysiological mechanism is yet unknown. It is still unclear how the ictal phases’ periods are determined? Why can any small trigger sometimes initiate the ictal phase, and sometimes, even bigger triggers cannot? Considering the brain as a dynamic system and proposing a complex system model for that as a migraineur or a healthy subject is a viable method. Here, the interaction between whole neurons is analyzed rather than individual neurons. This model is a complex system with a chaotic attractor. With parameter alternations, this attractor changes from one scroll to double scroll, representing a healthy or a migraineur brain. In the proposed system, the attractor’s borders are the regions where every small trigger can start the ictal phase, while the outer areas are the non-sensitive brain situations. We believe that MD and Chua’s systems have certain behavioral similarities. This study aimed to explain the function of MD and offer a theory that adequately describes its behavior. Finally, it has been tried to discuss some physiological evidences such as Migraine Generator Network (MGN), Cortical Spreading Depression (CSD), and the role of Serotonin and other substances in relation to the expressed hypotheses. This insight may propose newer methods for preventing or curing MD. Knowing the functioning of dynamic systems and finding similar behaviors with MD on the one hand, as well as linking physiological and pathophysiological evidence with a quantitative model can be very useful in better understanding, managing, and controlling the MD.

## Introduction

Migraine Disease (MD) is one of the primary headaches affecting a patient in every seven people worldwide [1]. This prevalent disease is a neurological disorder with many symptoms, including headache, nausea, vomiting, photophobia, phonophobia, osmophobia, etc. The most important and bothering symptom of the MD is the throbbing headache. The occurrence frequency of migraine ictal can be rare, weekly or even daily. MD is considered a chronic disorder with episodic manifestations [2]. The mechanism of MD is not entirely understood until now. Still, activation of some regions in the brainstem named Migraine Generator Network (MGN) and Cortical Spreading Depression (CSD) during a migraine ictal are accepted as the involved theories in the pathophysiology of MD. These theories have trouble in explaining the difference between a healthy brain and a migraineur brain in the inter-ictal phase. This disease has one normal and four abnormal phases, including pre-ictal, aura, ictal, and post-ictal phases. In some patients, just one or two disease phases are seen [3]. Two common subtypes of MD are migraine with aura (MA) and migraine without aura (MO), based on whether the migraineur experiences the aura phase. The diagnosis of MD is depended on the clinical symptoms that patients describe and the clinician’s opinion.

Some studies concentrate on classifying migraine and control subjects using features extracted from their electroencephalography (EEG) signal[1, 4, 5]. Some researchers have considered the brain a dynamic system that alternation of its parameters leads it to the MD state [6-8]. In 2003, Charles described migraine as a brain state. He stated that headache happens due to changes in the state of the brain. At the beginning of the ictal phase, a series of brain networks become active or inactive, and coordination between different parts of the brain is lost. Also, arousal decreases during the ictal phase, and symptoms such as fatigue, yawning, etc., occur. While awareness increases, the brain becomes more sensitive to light, smell, etc. In migraine, in addition to activating pain-sensing networks, physiological communication between different brain parts is disrupted. Various studies have reported changes in neuronal connections in the inter-ictal phase, as well as a decrease in theta wave of the Quantitative Electroencephalography (QEEG) in the preictal and ictal phases [9].

In 2013, Scheffer et *al*. considered a minimal model for migraine. They reported that stimulating a group of neurons by an input stimulus increases the intracellular potassium and glutamate, increasing the neurons’ excitability. Local neuronal activity and positive feedback further increase the excitability, and a small trigger initiates a contagious process in neurons called CSD. This study states that in every small area of the brain, there is a dynamic balance that results from the generation and decay of the pulses. When baseline excitability increases, the balance is lost, and the brain enters the tipping point. In this case, every small trigger initiates the ictal phase [8].

Dahlem et *al*. in 2013 considered migraine to be a dynamic disease and stated that when the headache starts, the brain comes out from the normal phase and enters a tipping point or bifurcation point and then enters the headache phase. They declared this stage the prodrome stage and stated that it could be identified with dynamical network biomarkers. Identifying this stage is vital since the prodrome stage, unlike the headache stage, is reversible to the normal phase, and perhaps there is a way to prevent the headache [7].

Also, Dahlem et *al*. in 2014 stated that when a brain enters a tipping point, even a small trigger can start the headache. Contrarily, when the brain is not in this area, even the items known to be the primary triggers of migraine headaches do not cause pain. They considered a path with one or two wells as two healthy and pain states. As the height between the two wells decreases, the brain enters the tipping point area, and each small trigger causes the headache to start [6].

In 2018, Bayani et *al*. expanded the model proposed by Scheffer et *al*. and considered a group of neurons for 3 different trigeminovascular, descending modulatory brainstem, and cortex units, then obtained 3 equations for neuronal activity. They also announced that the inter-ictal and the ictal stages are chaotic phases, and the pre-ictal is unstable and periodic [10].

According to previous studies, considering the brain as a dynamic system and then proposing a complex system model for that as a migraineur or a healthy subject can be a proper method. For this reason, it seems that by examining the behavior of dynamic systems that have similar performance to MD behavior, it is possible to gain a good understanding of MD performance. Also, this can be done to better understand the changes of the brain during the ictal phase for a better diagnosis and maybe better therapy.

### Hypothesis

MD is considered a chronic disease with episodic manifestations. Also, if the occurrence frequency of the headaches increases, chronification of the disease occurs. It results in chronic migraine with 15 times or more occurrences in a month for more than 3 months. It has also been believed that there is a pre-ictal stage before a migraine ictal. This stage can be a warn of the headache initiation with different symptoms, including behavior changes, hunger, fatigue, and etc. Despite some theories about the reason of the MD like CSD or MGN, the pathophysiological mechanism of the MD is not clearly understood yet.

There have been many computational models for MD during the 1970s, trying to explain the spreading depression. Some studies have also considered the Central Nervous System (CNS) role as the function of a complex dynamic system. As the parameters of the dynamic system change, the symptoms of the disease become apparent. Here a theoretical model is presented with the view of the complex dynamic system. This is used to explain the clinical signs and manifestation of MD. The dynamics of the individual neurons are not considered here, but the interaction between whole neurons is analyzed. A complex system with a chaotic strange attractor is regarded as the healthy brain model. This attractor is considered to change from a one scroll to a double scroll attractor with system parameter changes in migraineurs’ brain. It is assumed that the one scroll attractor can represent a healthy brain, whereas the double scroll one represents a migraineur brain. The size of the two scrolls can change by varying the model’s parameters to exhibit the Chronic Daily Migraine (CDM) or episodic migraine.

For a migraineur, some triggers such as neck pain, bright light, certain food, and other factors can sometimes start the ictal phase, but the same stimuli do not affect other days. It is believed that the brain becomes sensitive in some situations where any trigger can initiate the ictal phase. In the proposed system, the borders of the normal attractor are the regions where every small trigger can change the phase and, as a result, start the ictal phase. In the proposed model, the inner areas of the normal attractor are the non-sensitive brain situations.

### Computational Model

To choose a proper model for MD, different dynamical systems were considered. We aimed to find a system that has an attractor that represents a healthy brain and by parameter change, it is able to produce two attractors and can simulate the behavior of a migraineur brain. The assumed system should have a one scroll mode and a double scroll mode with two distinct scrolls representing the healthy brain and a migraineur brain. The alternation between the modes happens with a parameter change. One of the best-known systems having these features is Chua’s system. Chua is a non-linear and well-known dynamical model [11]. In 1983 Leon Chua developed a chaotic electronic circuit with 2 linear capacitors (C_1_, C_2_), a linear inductor (L), a linear resistor (R), and a voltage-controlled non-linear resistor (R_N_) known as Chua’s diode, which can produce chaotic behavior. This circuit is shown in figure 1. The Chua’s diode has 3 regions; a piecewise linear region with 2 unstable points. The driving points of Chua’s diode are shown in figure 2.

**Figure 1.**
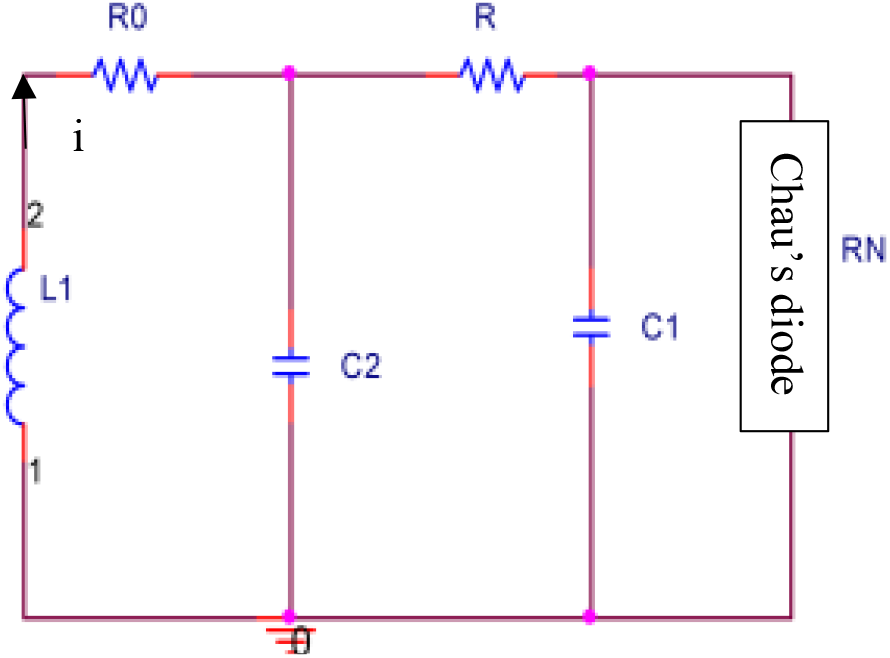
Chua’s circuit

**Figure 2.**
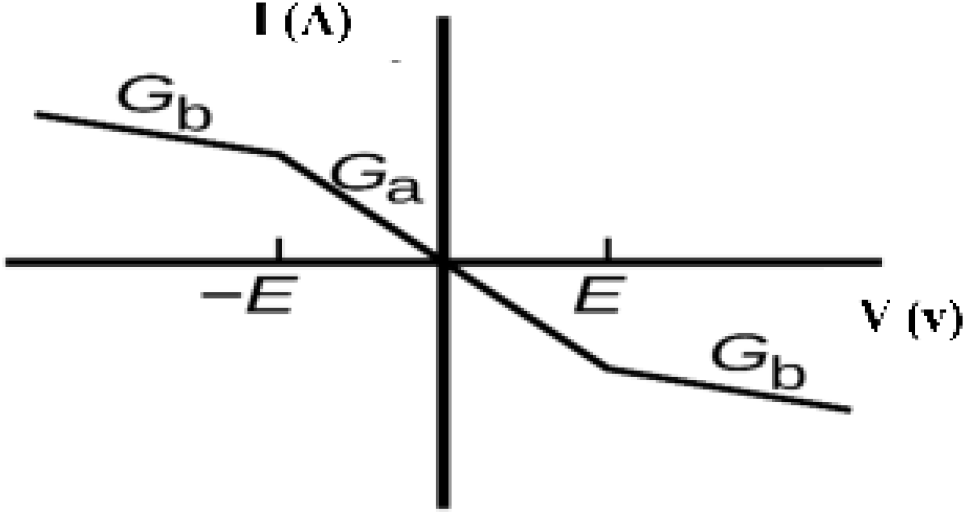
The driving point characteristics of the voltage controlled resistor

The circuit’s equations that are derived by applying Kirchoff’s laws on nodes and loops are given as:

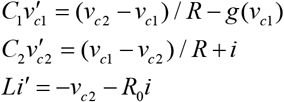

Where C_1_ and C_2_ are the capacities of the capacitors, L is the inductance of the inductor, R is the resistance of the resistor, and the g(x) is the three-segment piecewise linear characteristics of the Chua’s diode, which is shown in figure 2 and mentioned here:

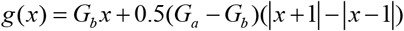

G_a_ is the slope of the inner region of the non-linear resistor, and G_b_ is the slope of the outer part. Then the variables of these equations are replaced with x_1_=v_1_, x_2_=v_2_, x_3_=Ri, and the parameters with

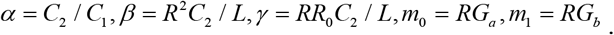

Chua’s system equations are achieved as:

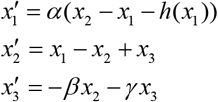

Where

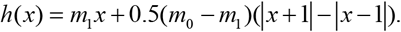

For different values of parameters, this attractor changes from a spiral attractor to a double scroll one. The spiral attractor can be a representation of a healthy subject, whereas the double scroll one can be a representation of a migraineur brain. Every region can show the ictal or inter-ictal phases, and the lines connecting the two areas show the pre-ictal and post-ictal phases.

### The similarities between MD and Chua’s system

A healthy brain that does not experiences a migraine ictal has just a normal phase. This mode is the one scroll or spiral mode of Chua’s attractor, as shown in figure 3.

**Figure 3.**
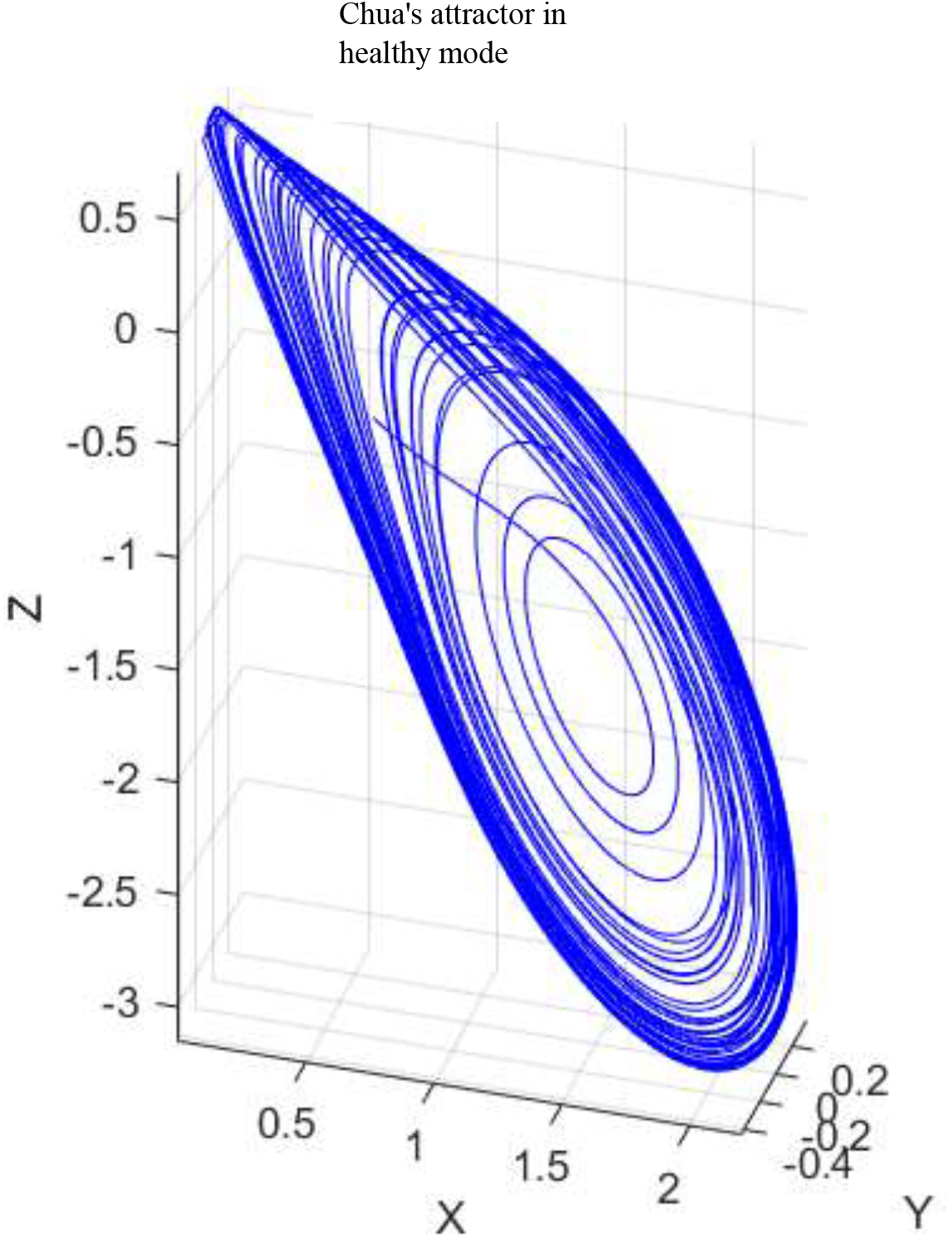
Chua’s system in the spiral mode representing a healthy brain

The value of the parameters for this mode is as:

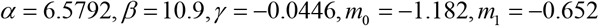

Also, the shape of the x(t) in this region is shown in figure 6. a.

A healthy brain has just the normal phase that is headache-free. In the proposed model, the change of the parameter alpha can change the healthy brain to a migraineur. By increasing the alpha parameter, the attractor, which has two scrolls indicating two phases of the brain of a migraineur, is obtained: the interictal or headache-free phase and the ictal or headache phase are connected with some lines to each other. Figure 4 is a representation of the double scroll attractor. The value of the parameters are as:

**Figure 4.**
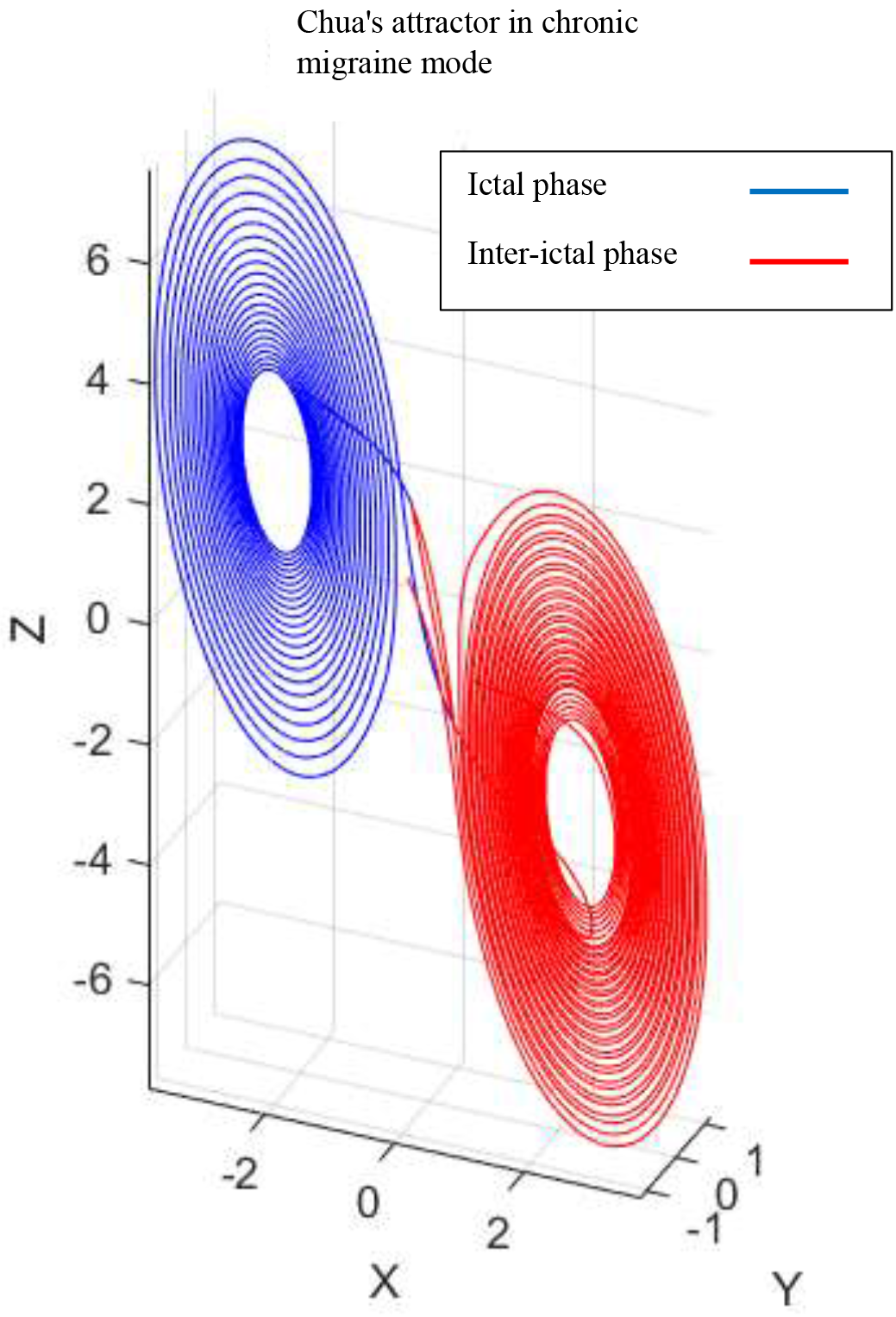
double scroll Chua’s attractor representing a chronic migraineur brain

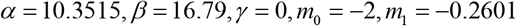

Every region of the attractor represents the ictal and inter-ictal phases. Suppose the size of one of the regions is smaller than the other, as it is shown in figure 5. In that case, the small region can represent the ictal phase for the situation of the episodic migraine in which headache occurs in lower frequency. But if the size of the two areas is the same, it can represent the brain of a chronic daily migraineur in which the migraineurs experience headaches every 15 days of a month or more. For the parameter values as:

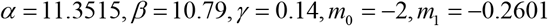

The Chua’s attractor becomes as figure 5 representing the brain of an episodic migraineur.

**Figure 5.**
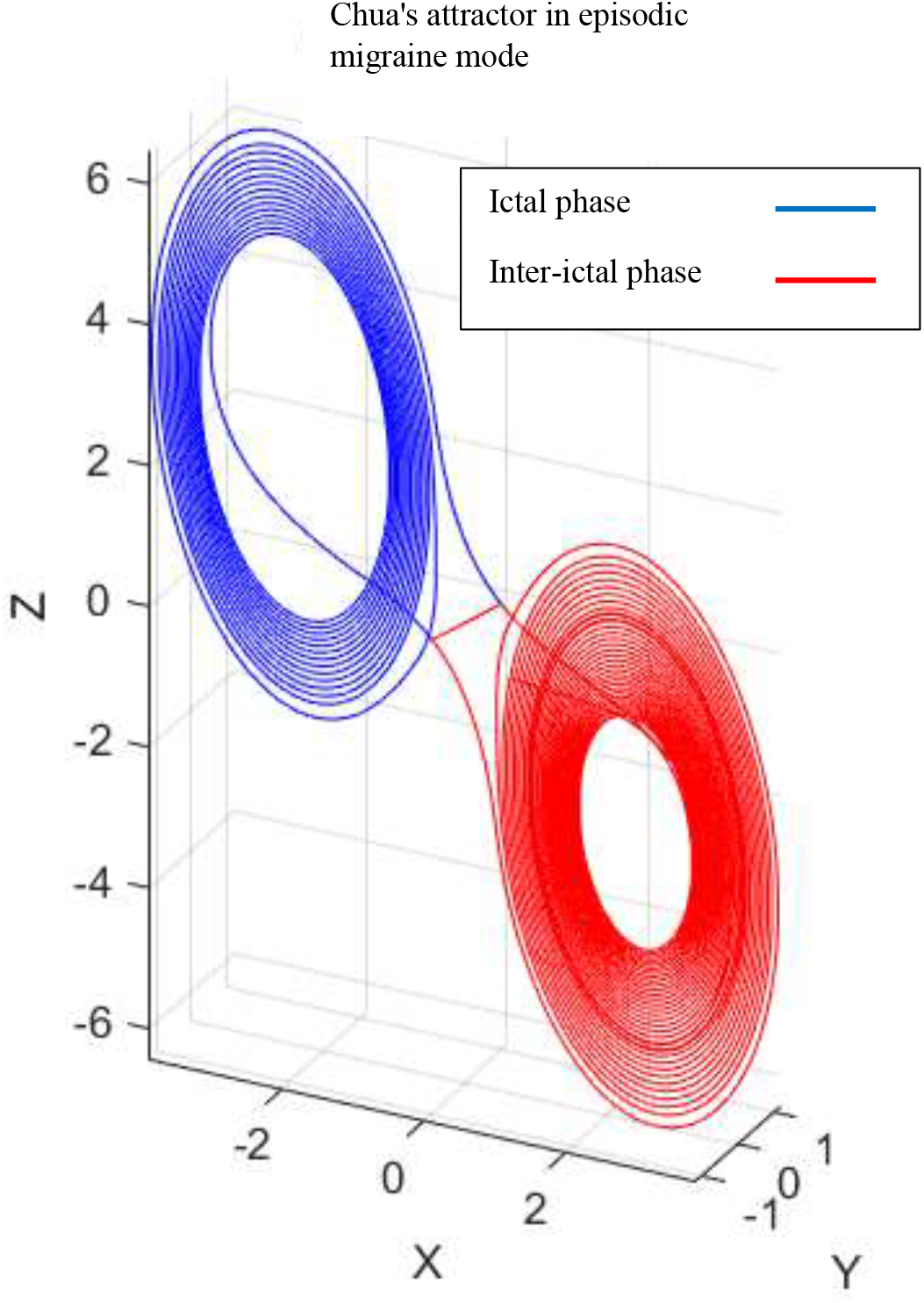
double scroll Chua’s attractor representing an episodic migraineur brain

**Figure 6.**
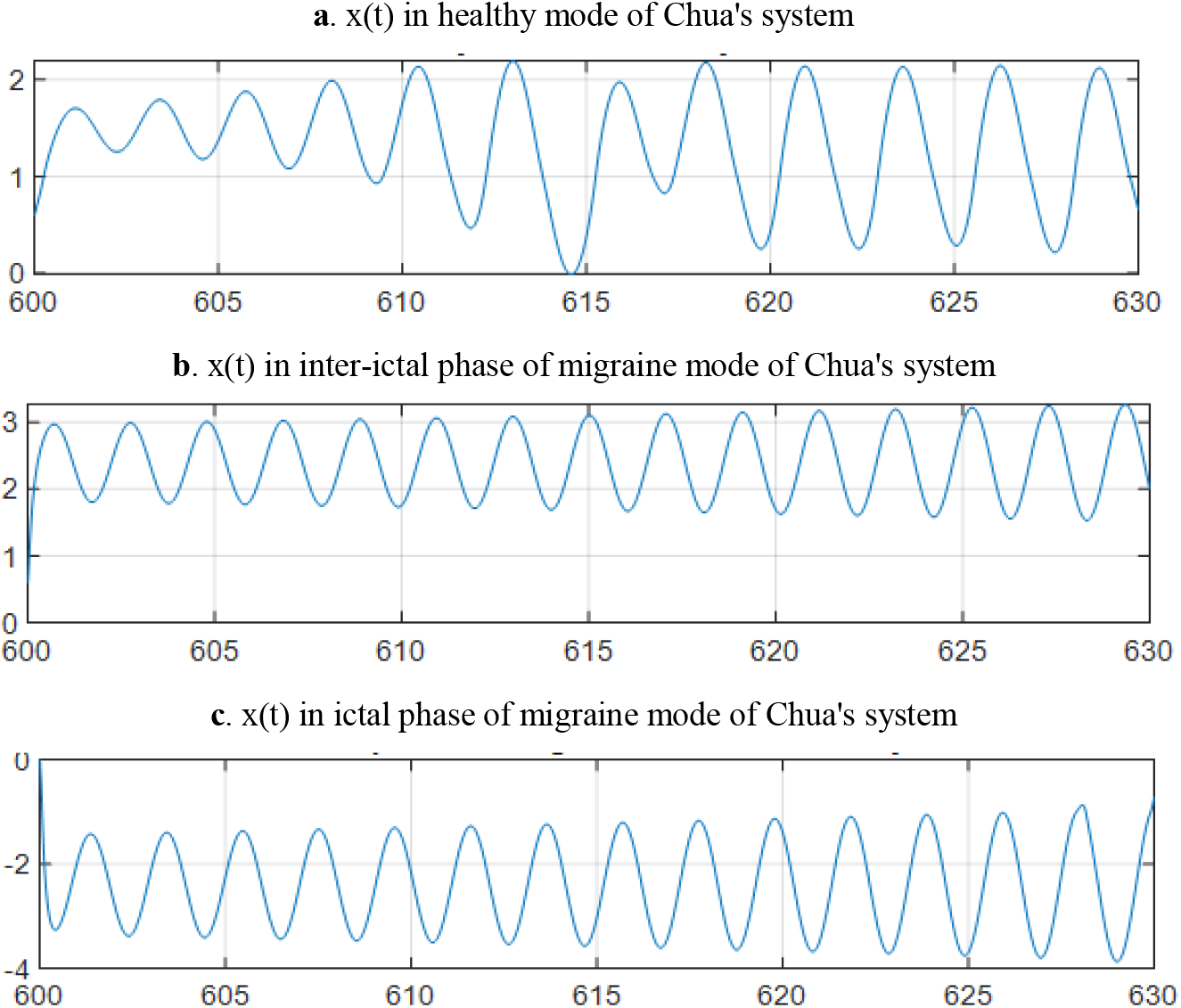
a. Shape of the x(t) in the healthy mode. b. Shape of the x(t) in inter-ictal phase of the migraine mode. c. Shape of the x(t) in ictal phase of the migraine mode.

The x(t) representation in healthy mode, ictal, and inter-ictal phases is shown in figures 6.a, 6. b, and 6. c. As shown in figure 6, the behaviors of x (t) in the 3 states of the healthy mode, inter-ictal phase, and ictal phase of the migraine does not seem to make a significant difference. If we compare the EEG of these 3 groups, no tangible difference in the appearance of the EEG in the time domain of these 3 groups is seen. But in general, these behaviors indicate whether or not there is the MD. As a result, we can say that what happens in epilepsy does not occur in MD. In epilepsy, these differences are seen in the EEG because of the synchronization. However, in MD, there is no tangible difference in the EEG’s appearance. In this figure, also there is no substantial difference between the behavior of the three groups. Still, they originate from different phases and modes of the Chua’s attractor and result in other states. Also, by comparing the x(t)’s energy level in the different modes of Chua’s system, differences in the energy levels is seen. The energy level was lower in Chua’s spiral mode (here considered as healthy mode), than the double scroll mode (considered as migraine disease mode in this study). By calculating the energy levels of the EEG records of migraineurs and healthy controls in the inter-ictal phase using the publically available dataset provided by Carnegie Mellon University [12], this higher level of mean energy was also seen in migraineurs’ brains than healthy controls. These values are expressed in Table 1.

**Table 1.** Comparison of the the different modes of the Chua’s system’ energy levels and EEG records of the migraineurs and healthy controls

| Energy level in one scroll mode of chua's system (healthy mode) | Energy level in double scroll mode of chua's system (disease mode) |
| --- | --- |
| 1.9574e+06 | 5.8743e+06 |
| Mean energy level in healthy controls' EEG record (mean of all channels) | Mean energy level in migraineurs' EEG record in rest (mean of all channels) |
| 15839 | 22957 |

It is also mentioned by the researches that higher glutamate level is seen in migraineurs’ brains than healthy controls [13]. Since glutamate has a significant role in brain energy metabolism and neuron excitation [14], maybe the reason for the higher energy level is the higher glutamate level in migraineurs’ brains.

It is belived by migraineurs that sometimes any small trigger can initiate migraine ictal, while in other situations even bigger triggers cannot start the ictal phase. In order to test if this situation can be seen in the proposed model, a point is chosen in the borders of the inter-ictal phase, as seen in figure 7, little noise is added to test if the point enters the ictal phase or not. This point enters the lines between two phases which is a way to the ictal phase. This noise can simulate the small trigger which initiates a headache. Also, the lines between the two phases can be considered pre-ictal and post-ictal phases of the migraine. Another point is also chosen in the inter-ictal phase inner area; then, the same noise is added to the system. Since the trajectory is in the inter-ictal phase and far from the borders, the same noise cannot initiate the headache and take the trajectory to the ictal phase.

**Figure 7.**
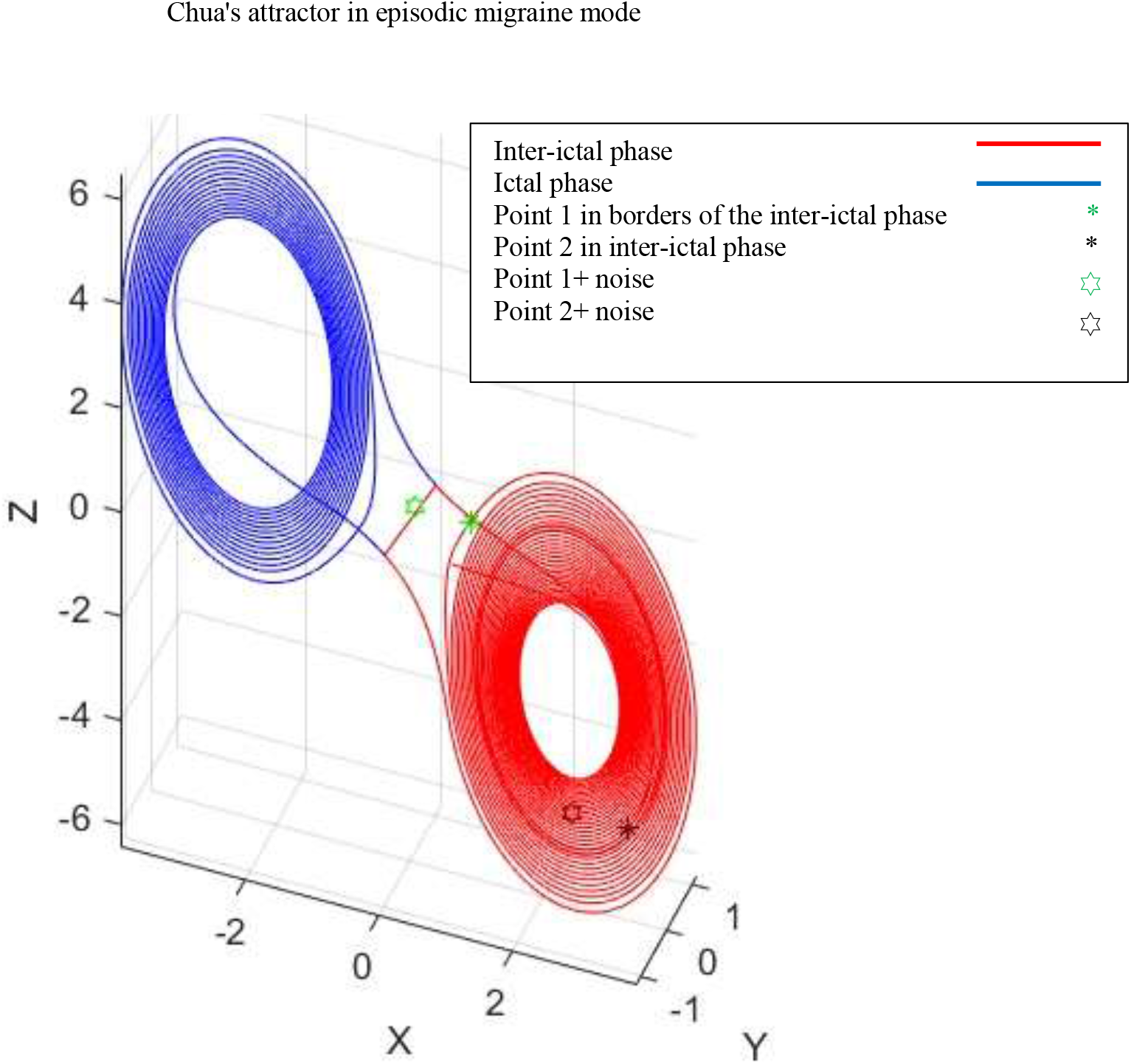
A point in the border of the inter-ictal phase entering ictal phase

## Discussion

Headaches are among the common disorders that 90% of people experience at least one time in their life. Episodic migraine is a common primary headache type that affects 15% of women and 6% of men worldwide [15]. 20% of the migraineurs experience migraine with aura [3]. The prevalence of migraine increases from childhood until 40 years old, and after that, this prevalence declines [16]. For some migraineurs, the frequency of the occurrence of the headaches increases where these people experience migraine without aura 15 times a month. If it lasts for more than three months, this disorder is known as CDM [17]. MD has affected many people worldwide, especially younger people, and has prevented the patients from their everyday lives. 53% of migraineurs report a need for bed rest, and one-third miss their school or job for one day in a year [17]. There isn’t any specific cure to prevent the occurrence of this disease.

Despite the high prevalence of MD and the disabling, which is the result of the severe pain during the migraine ictal phase, the reason behind the disease is not clearly understood yet. Although many studies have focused on the disease pathophysiology [18-20], some questions about MD have remained unanswered. What is the real reason behind the occurrence of the MD? Medical texts cite two mechanisms for MD, but it is not yet clear how these mechanisms are associated with headaches and why new manners occur? How are the periods of migraine ictal determined? Why can any small trigger sometimes initiate a migraine headache and even bigger triggers cannot start the headache in other situations? What is the reason that headaches get worse or weak? What is the nature of the pre-ictal and the post-ictal phases of migraine headaches?

It seems that the study of individual neurons and changes in MD is not very appropriate for assessing and recognizing the function of the disease. We think that if we have a global view of the neuronal areas, and the function of the area is considered, a better understanding can be obtained. Given the knowledge about the performance of the dynamic systems and various studies that have compared the performance of the CNS with that of dynamic systems, it can be concluded that having a view of dynamic systems for analyzing brain function may lead to a better understanding of MD. Therefore, we hypothesize that different neuronal regions from the brain’s normal behavior can be considered an attractor that the system response remains in this healthy attractor. In the event of MD, a new attractor is added to the responses of different areas of CNS, which can indicate the disease attractor. Because entering the abnormal area distorts the function of the brain, this distortion is interpreted as pain in the brain and forms a classic migraine pain. Although there aren’t any pain detection sensors in large parts of the brain, severe migraine pain can be caused by mismatched information of the different areas of the brain together. The best example of this mechanism is the take-off moment of the airplanes. Although the vestibular system senses high acceleration, the visual and auditory systems do not perceive this acceleration, and this mismatching of the information causes headaches during take-off. According to this theory, some neuronal regions referred to as MGN in medical texts produce migraine attractor regions. When the number of these neuronal regions is more significant, the mismatching rate is higher and severe headaches are felt.

The model of a new attractor’s creation is closer to reality. In fact, there is no headache when the brain is in the first attractor, and a headache occurs when the second attractor is created and the brain enters this attractor. It can be said that the creation of the second attractor occurs due to the activation of MGN or CSD as a result of changes in the levels of substances such as serotonin, glutamate or potassium. Then the brain enters the second attractor. After creation of the second attractor, when brain enters this attractor, it has been observed that the brain function energy (EEG behavior) increases, resulting in elevation of the cell metabolism, increased number of action potentials and enhancement of ATP consumption. The EEG records of migraineurs’ brains also show an increase in energy level as well as more spikes than healthy controls. This ATP consumption cannot continue indefinitely and has to be reduced to some extent and return to the previous state. Creation of the second attractor occurs by activation of MGN and CSD, then the increased level of glutamate leads to entry in the second attractor. Then an increase in energy level and ATP consumption occurs. After a while, the brain is forced to leave this second attractor and return to the normal state. Also, it is recently demonstrated that ATP sensitive potassium (KATP) channels are opened during migraine attacks [21]. In this study it is assumed that the energy level in one part of the brain increases suddenly due to an increase in glutamate level, and other parts of the brain cannot adapt to these energy alternations and an information mismatching occurs.

One of the standard dynamic systems that exhibit complex behavior is the Chua’s system, where a remarkable resemblance between the behavior of this system and the MD is seen. According to the proposed theory, when a person does not feel a migraine headache, the levels of the glutamate and serotonin are altered a little, then the brain compensates these alternations and most of the MGN neuronal areas in the attractor are in normal function. Then the mismatching rate in different brain areas is low, and the headache does not occur. When the alternation of the glutamate and serotonin is high, MGN neuronal states are activated, brain enters the second attractor, energy increases and mismatching of the information occurs then the headache is felt. Since only the normal attractor is formed in healthy people then, in migraine patients, the attractor related to the headache area is developed gradually. Therefore the growth of the disease can be justified by enlarging the ictal area of the attractor. By comparing the size of the migraine ictal attractor to the inter-ictal one, the length and shortness of the headache, as well as the frequency of migraine headaches, are justified.

In some cases, patients become severely sensitive to stimuli depending on the proximity of the trajectory to the branch area, which is the passage to the ictal attractor. The farther the trajectory is from the branch, the greater the stimulus needed to enter the ictal area. In contrast, the closer the trajectory is to the branch area, the trajectory enters the migraine ictal area with a smaller stimulus. According to the drawn trajectory and the phase space of figure 7, the attractor transition areas can be medically considered preictal and post-ictal areas. In the pre-ictal area (the transition path from an inter-ictal attractor to the ictal attractor), it gradually deviates from normal function. The headache rate gradually increases. The transition time can be short or long between different people. In the post-ictal area (the transition path from the migraine ictal attractor to the inter-ictal attractor), we move away from the migraine-related areas with severe headaches and approach the inter-ictal attractor, the amount of pain gradually decreases.

## Conclusion

The anatomical variations between the brains of healthy people and migraineurs, which may be caused by genetic factors or other circumstances, are thought to be the reason of the creation of the second attractor, which is the attractor of the ictal phase of the migraine. CSD and MGN can occur in migraineurs’ brains leading to the ictal phase of the migraine. Then the difference between the structure of the migraineurs’ brains has given rise to a second attractor.

The reason of the switch from inter-ictal phase (first attractor) to the ictal phase (second attractor) in migraineurs is a decrease in serotonin level when brain is in the borders of the first attractor. If serotonin level decreases in migraineurs’ brains for any reason, which is normal in daily brain behaviors, and also the brain is in the borders of the attractor, it enters the ictal phase of the migraine (second attractor). However, if the brain is not at the borders of the attractor, the decrease in serotonin level is compensated by brain and does not cause the brain to enter the second attractor. This is why sometimes every small stimulus triggers a headache, but in other cases, even larger stimuli have no effect.

The switching factor from ictal phase to inter-ictal phase in migraineurs, is the disruption of coordination in the structures formed in the brain of migraineurs. The structures which are a result of CSD and MGN activation. These structures are disconnected when the neurons in this region do not generate needed action potentials. When the ATP intake of the neurons increases due to increased energy level, a lack of ATP and energy occurs, then the neurons do not generate action potentials for resumption of the path, so the headaches are ceased. In other words, the energy level in the brains’ of the migraineurs is higher, after a while, the body cannot provide this higher energy level, and as a result, the neurons in the area involved in the headache are deactivated and the second attractor is interrupted. The interruption of the second attractor leads in reversal to the first attractor, then the headache is stopped.

The cause of the headaches is mismatching of the energy levels occurring in migraineurs’ brains. The brain of a migraineur is initially adapted to a lower energy level. Suddenly, the energy level of a part of the brain increases then the brain cannot adapt to this higher energy level and the headache occurs.

According to the above mentioned items, since ATP plays a key role in the migraine cycle, high fat and high calorie foods keep the second attractor on standby. In this case, a brief stimulus starts the headache phase. When migraineurs eat high-fat, high-calorie foods, they are more likely to enter the second attractor, and more likely to have headaches [22]. It can also be said that if migraineurs consume high-calorie and high-fat foods during the headache, their headaches last longer. It is recommended that, migraineurs eat less fatty and low-calorie foods, especially when they feel close to the attack phase.

In this study, it has been tried to explain the disease function and offer a theory that reasonably justifies the behavior of the MD. This explanation may propose newer methods for preventing or curing the MD. To better understand MD to control it and shrink the areas involved in this disease, it is better to know the dynamic systems better. It may help prevent the formation of migraine ictal attractor or even make the migraine ictal phase attractor smaller even after it has been formed. A series of electrical stimuli when the headache starts can take us back from the migraine ictal area to the inter-ictal area, which should be further studied.

